## Supplementary Information for "Consistent Tumorigenesis with Self-Assembled Hydrogels Enables High-powered Murine Cancer Studies"

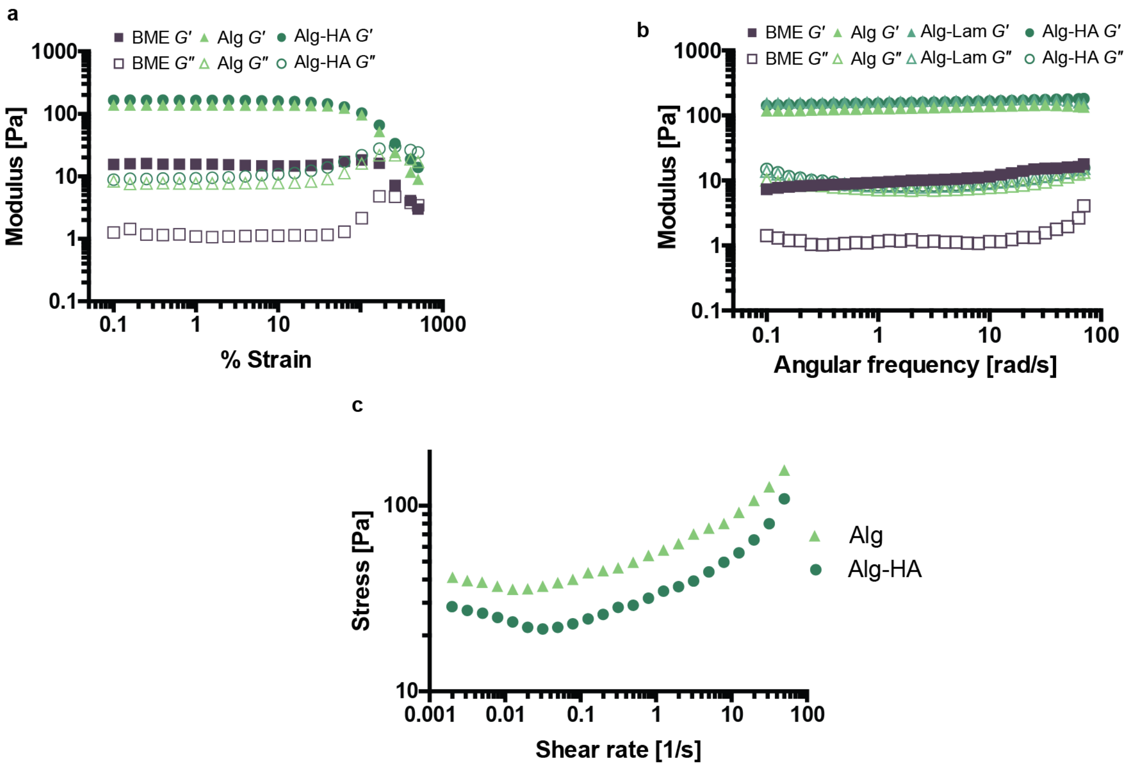


**Supplementary Figure 1: a,** Amplitude sweep (w=10 rad/s) for all formulations at 37°C. All hydrogel formulations show a similar stiffness, a similar solid-like response, and a similar range of linear viscoelasticity across the measured strains. BME is less stiff, but yields at a similar strain. **b,** Frequency sweep (strain=1%) for all formulations at 37°C. All hydrogel formulations show a similar stiffness, a similar solid-like response, and a similar range of linear viscoelasticity across the measured frequencies. BME is less stiff, but demonstrates a similar solid-like response. **c,** Flow sweep for Alginate and Alg-HA formulations at 25°C. Both formulations show a yield stress of approximately 20 Pa.

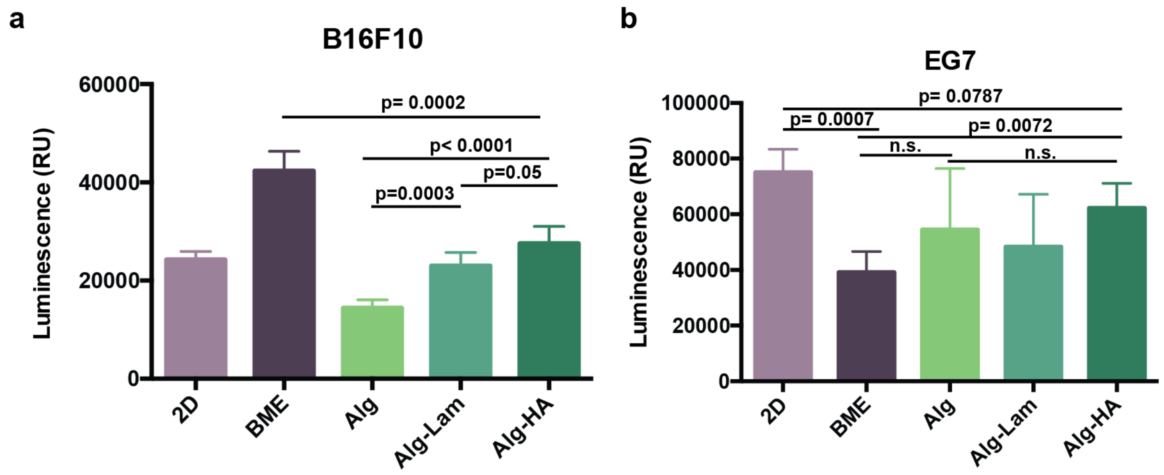


**Supplementary Figure 2: a,** Relative viability of B16F10 cells in various formulations after 1 day in culture after seeding 5,000 cells. Luminescent signal is linear to the cell number. **b,** Relative viability of of EG7 cells after seeding 10,000 cells after 1 day in culture in various formulations. Error is reported as non-significant (n.s.) if $p < 0.05$.

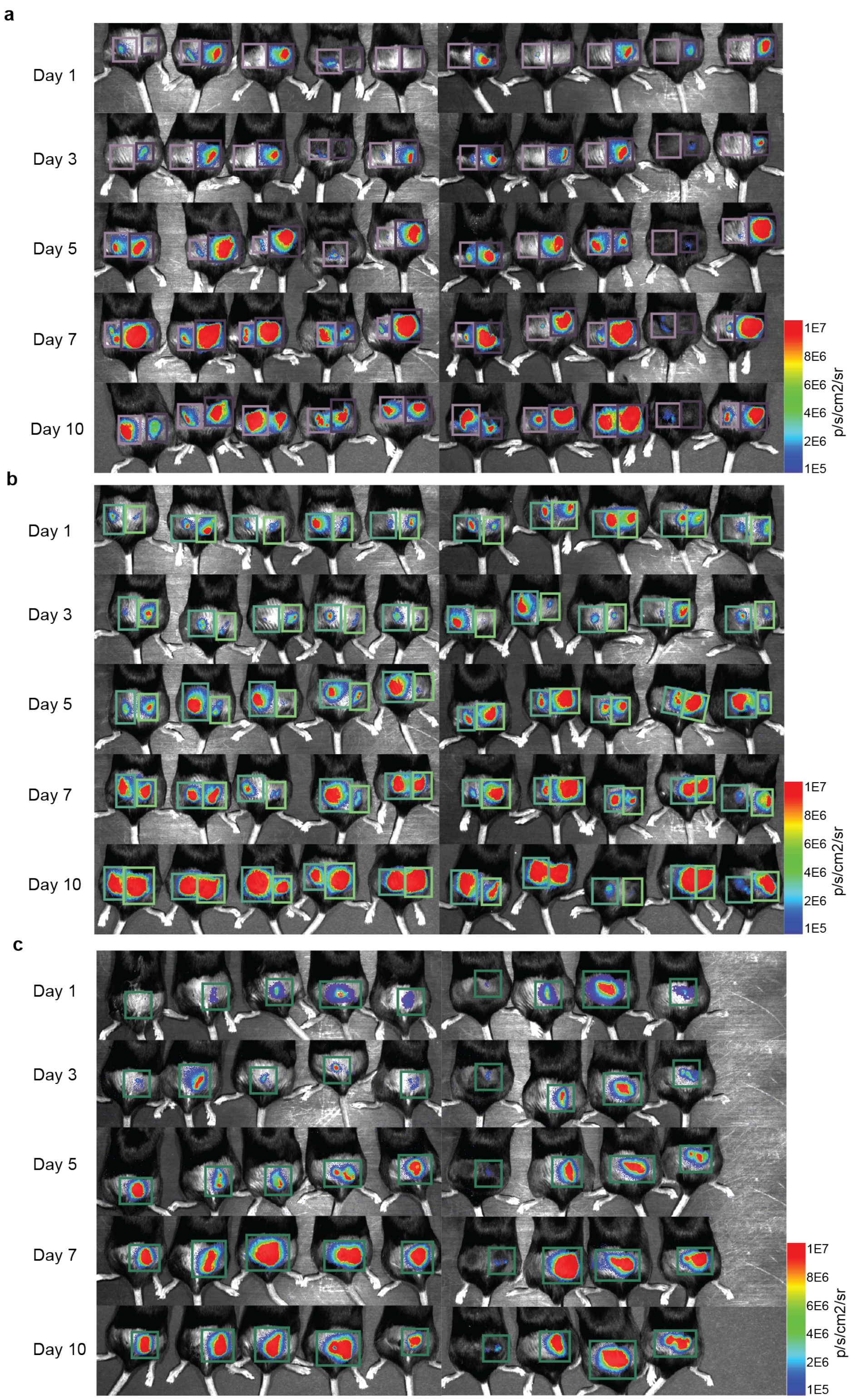


**Supplementary Figure 3:** *In vivo* imaging over time of mice with luminescent tumors forming. Tumors were innoculated with cells dispersed in saline, BME, Alg, Alg-Lam and Alg-HA. **a,** Saline (left, light purple) and BME (right, dark purple) groups. **b,** Alginate-Laminin (left, medium green) and Alginate (right, light green) groups. **c,** Alginate-Hyaluronic Acid (center, dark green) group.

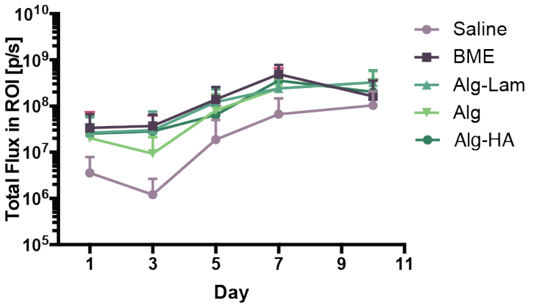


**Supplementary Figure 4:** The average total flux in region of interest surrounding the tumor of interest and standard deviation from in vivo imaging on various days directly after inoculation for all experimental groups.

Gompertz Model: $X(t) = K \exp\left(\log\left(\frac{X(0)}{K}\right)\exp(-\alpha t)\right)$

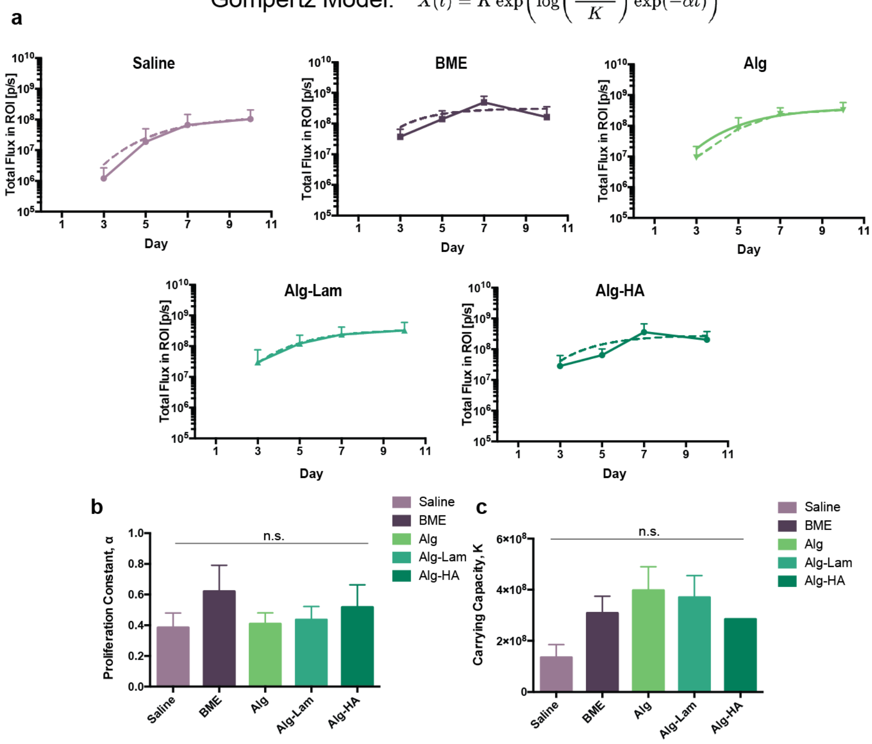


**Supplementary Figure 5:** *In vivo* imaging fit to Gompertz model. Parameter $\alpha$ that represents the proliferation constant, parameter $K$ that represents the carrying capacity, *X(0)* represents the initial size, *t* represents time (days). **a,** Tumor luminescence fits to Gompertz equation after initial delivery phase (starting on Day 3, see methods for details). **b,** Parameter $\alpha$ in the Gompertz model and standard error calculated from the fit for all groups. **c,** Parameter $K$ from the Gompertz model and standard error calculated from the fit for all groups.

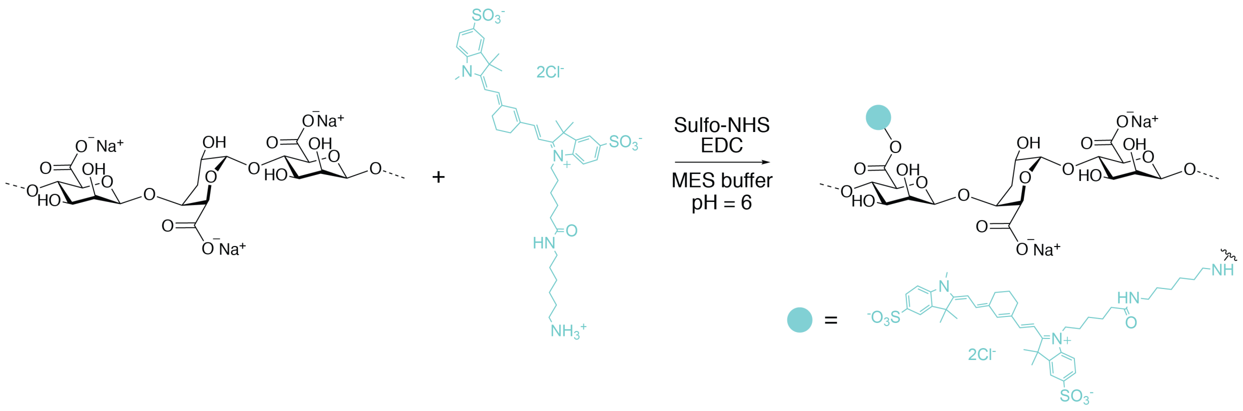


**Supplementary Figure 6:** Cy7 dye conjugation to alginate using carbodiimide chemistry.

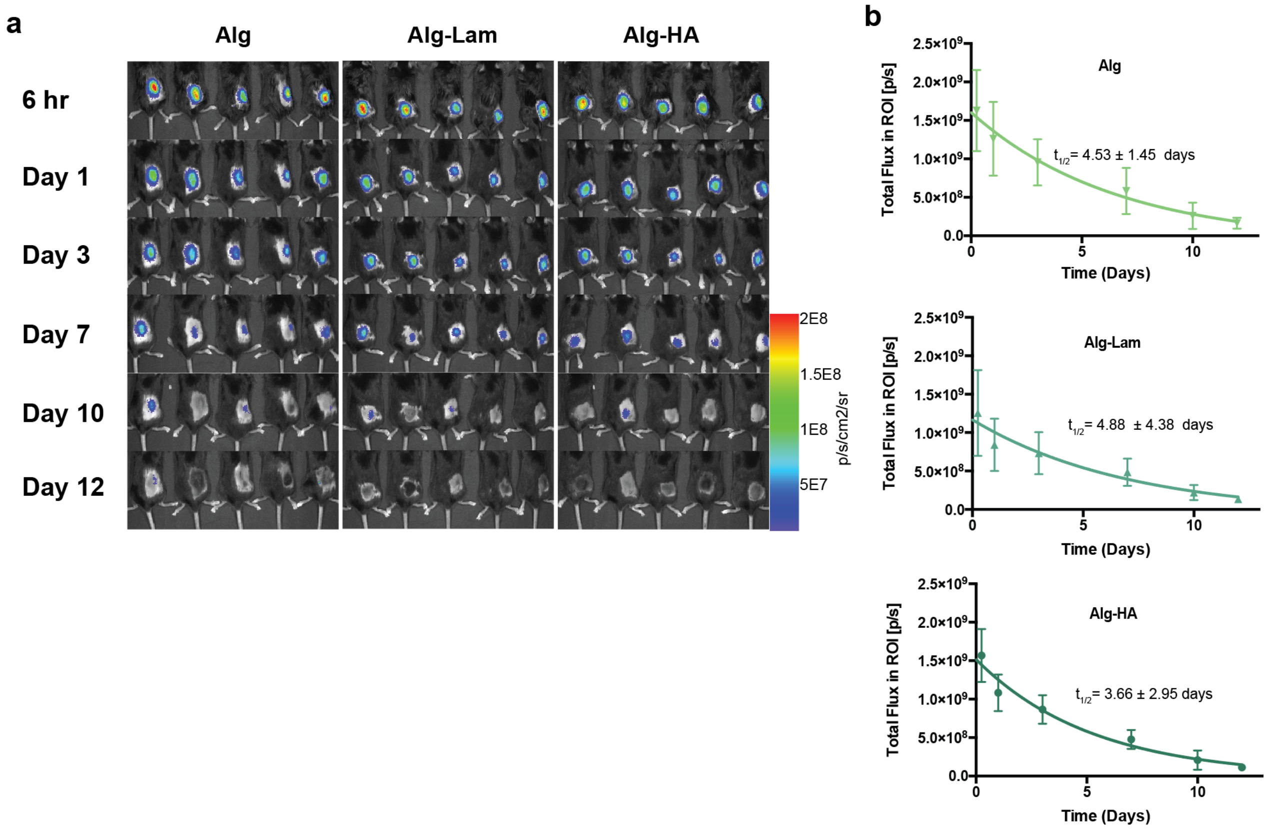


**Supplementary Figure 7: a,** *In vivo* degradation of fluorescent alginate measured using an *in vivo* imaging system for all hydrogel groups. B16F10 cells were encapsulated and co-injected in all groups as in previous studies. **b,** Average total flux and standard deviation from the region of interest surrounding the injection over time. A one-phase exponential decay was fitted to each curve and a half-life was computed for each group as shown on the graph.

**Supplementary Table 1:** P-values calculated with two-sided t-tests from analysis of tumor areas for all groups (Fig 2).

| Group Pair | Day 10 | Day 13 | Day 15 |
|---|---|---|---|
| Saline, BME | 0.0002 | 0.0011 | 0.0045 |
| Saline, Alg | <0.0001 | 0.0239 | 0.2778 |
| Saline, Alg-Lam | <0.0001 | 0.0014 | 0.0290 |
| Saline, Alg-HA | <0.0001 | 0.0015 | 0.0027 |
| BME, Alg | 0.5848 | 0.0163 | 0.0076 |
| BME, Alg-Lam | 0.9994 | 0.2423 | 0.0978 |
| BME, Alg-HA | 0.2800 | 0.2251 | 0.3943 |
| Alg, Alg-Lam | 1.0000 | 0.0282 | 0.0501 |
| Alg, Alg-HA | 1.0000 | 0.0113 | 0.0003 |
| Alg-Lam, Alg-HA | 0.1637 | 0.9485 | 0.1133 |

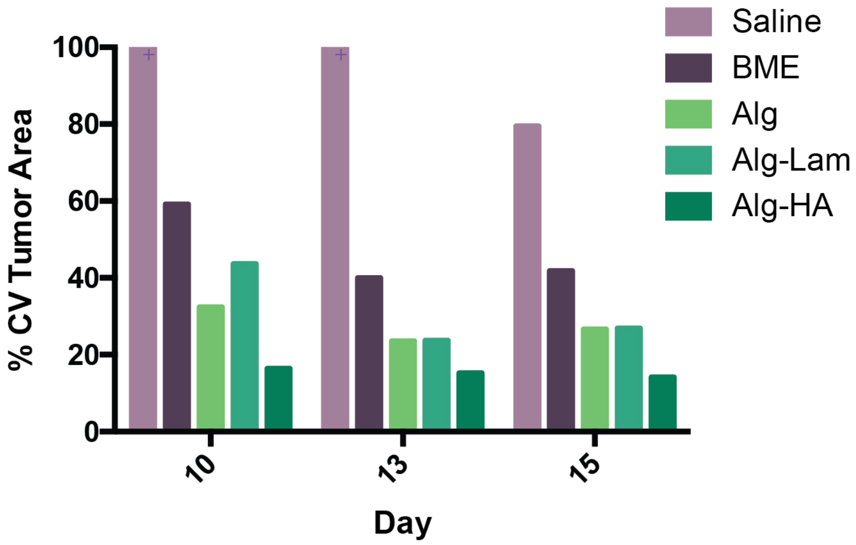


**Supplementary Figure 8:** %CV of tumor area for all groups: saline, BME, Alg, Alg-Lam and Alg-HA. Same data as Fig 2a plotted on one graph to show direct comparisons. Saline exceeded an 100 %CV on Days 10 and 13.

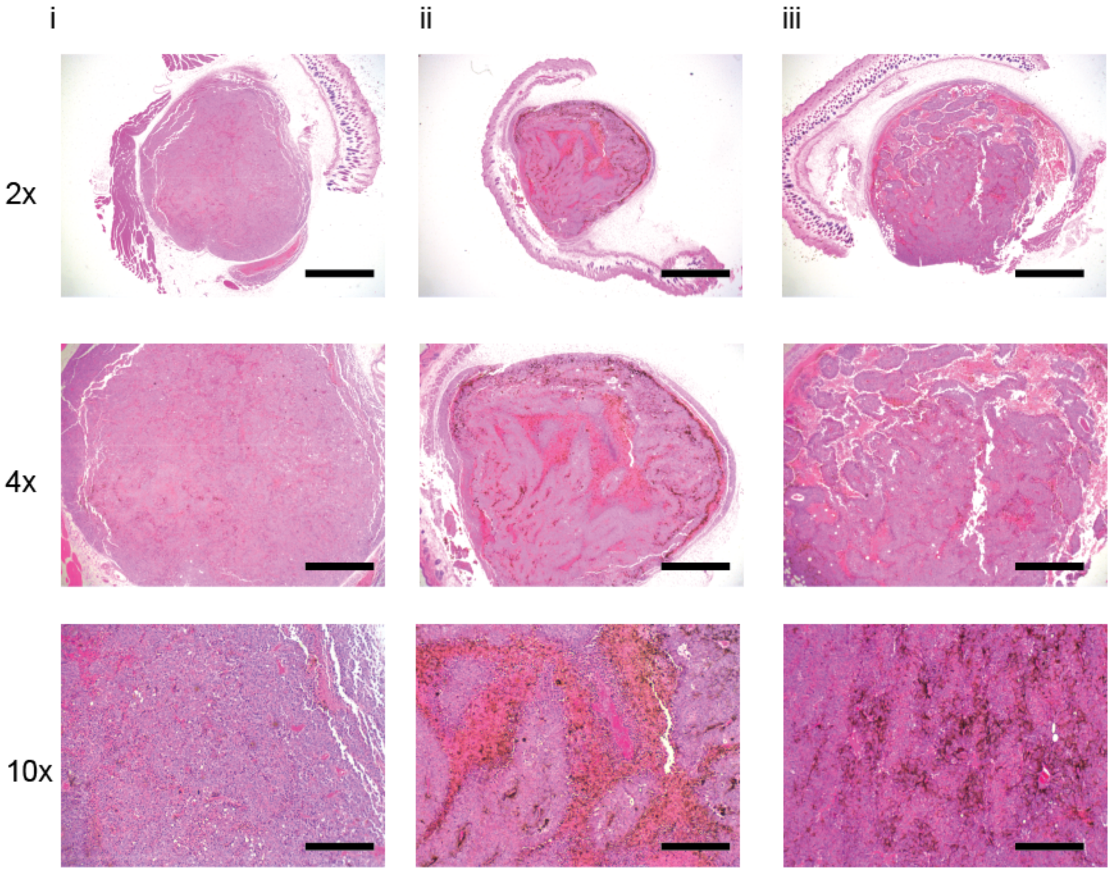


**Supplementary Figure 9:** Additional histology images with Hemotoxylin and Eosin staining from Day 15 for tumors formed in saline. Columns represent a single tumor, and rows represent a single magnification. Saline tumors were smaller than other groups. 2x magnification the scale bars represent 2 mm. 4x magnification the scale bars represent 1 mm. 10x magnification the scale bars represent 400 um.

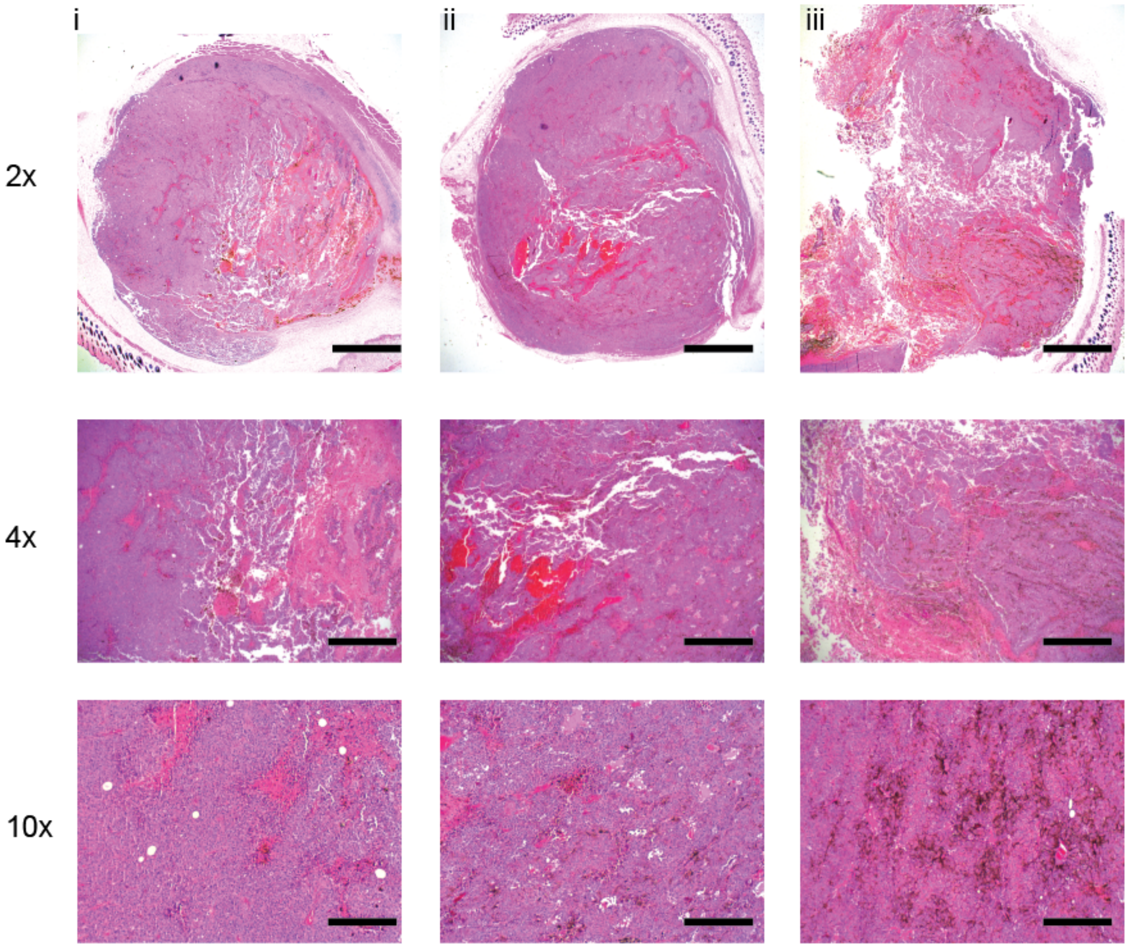


**Supplementary Figure 10:** Additional histology images with Hemotoxylin and Eosin staining from Day 15 for tumors formed in BME. Columns represent a single tumor, and rows represent a single magnification. BME formed the largest tumors of all groups. 2x magnification the scale bars represent 2 mm. 4x magnification the scale bars represent 1 mm. 10x magnification the scale bars represent 400 um.

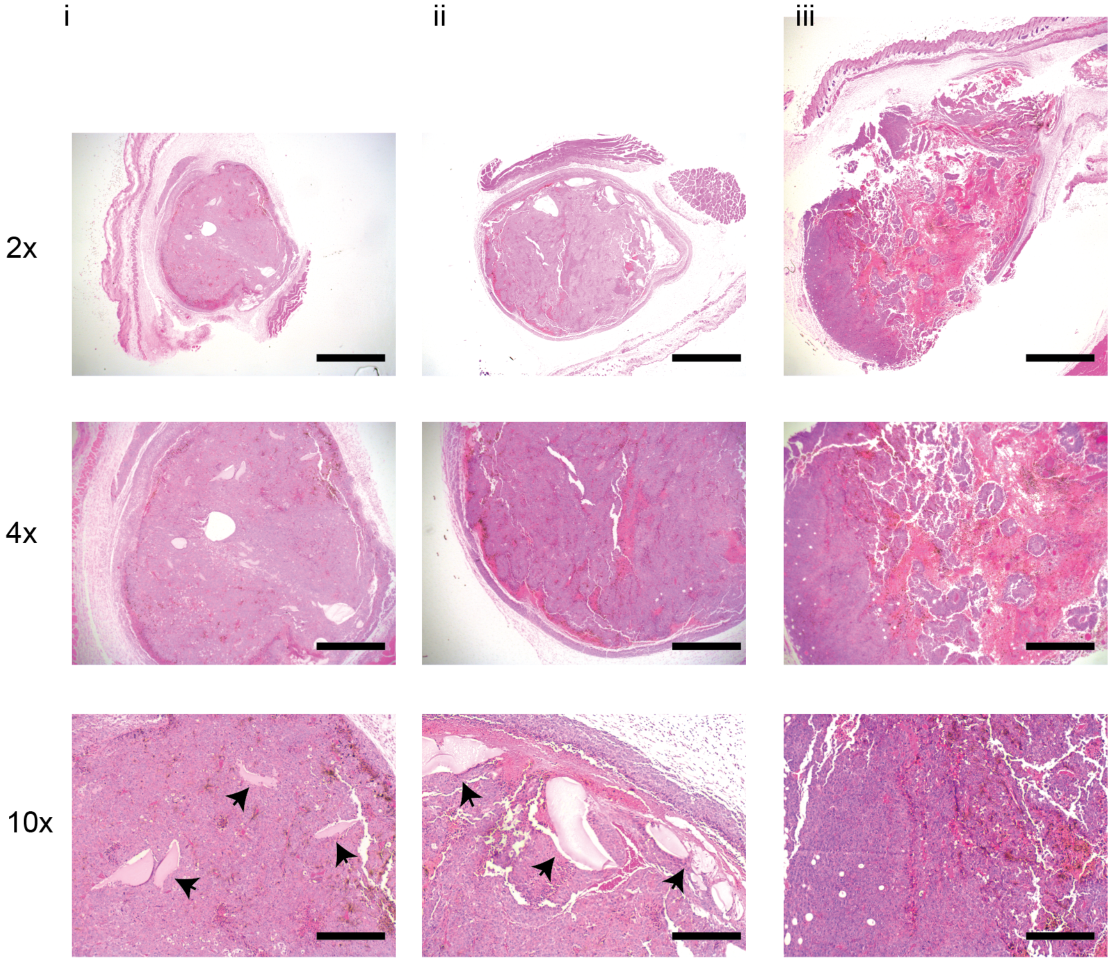


**Supplementary Figure 11:** Additional histology images with Hemotoxylin and Eosin staining from Day 15 for tumors formed in Alginate. Microscopic examination by a pathologist confirmed that the melanomas demonstrated identical histomorphologic features across all formulations. There is amorphous, pale eosinophilic material additionally present in some tumors formed in gels that may represent residual gel material. This material is designated by black arrows. Upon evaluation by a blinded pathologist, these tumors do not appear morphologically different than tumors formed in BME and saline. These tumors formed in alginate were smaller than the other gel groups. Columns represent a single tumor, and rows represent a single magnification. 2x magnification the scale bars represent 2 mm. 4x magnification the scale bars represent 1 mm. 10x magnification the scale bars represent 400 um.

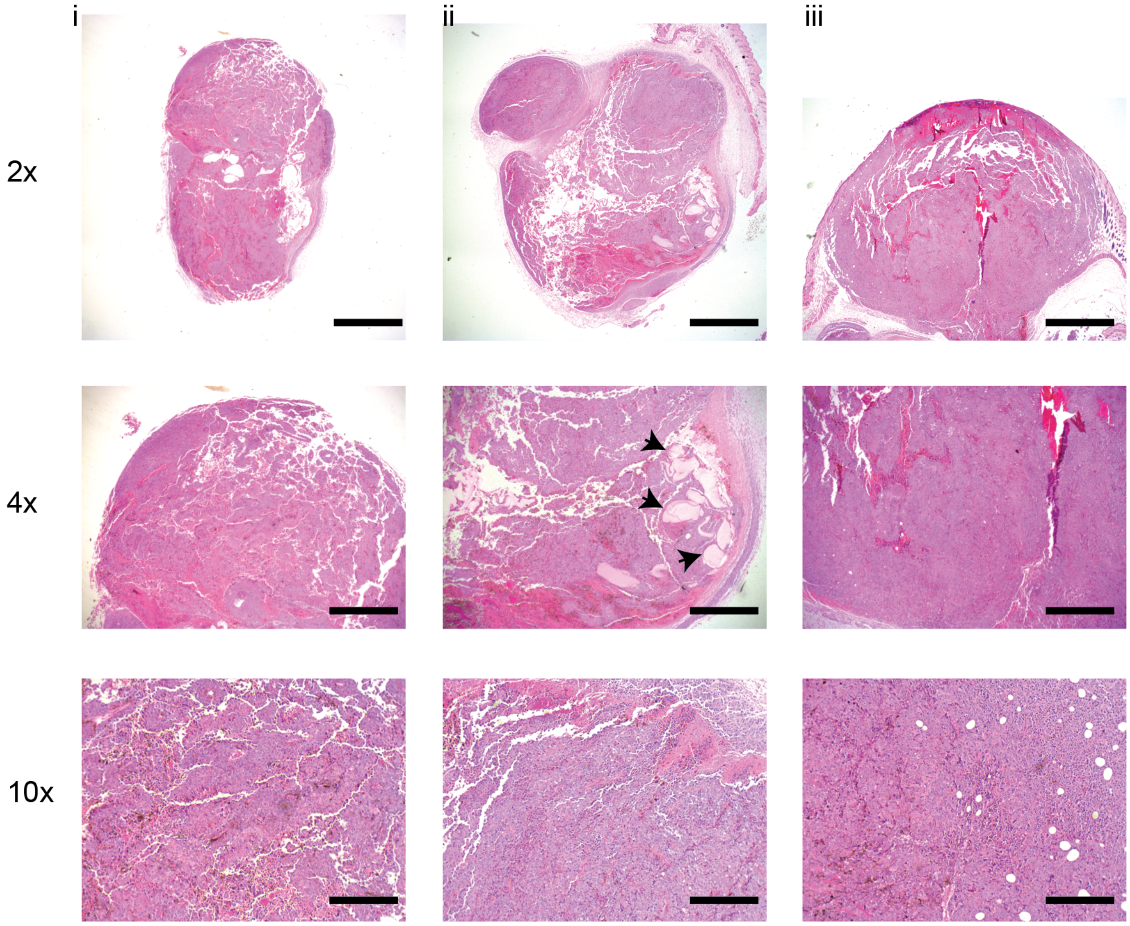


**Supplementary Figure 12:** Additional histology images with Hemotoxylin and Eosin staining from Day 15 for tumors formed in Alginate-Laminin. There is amorphous, pale eosinophilic material additionally present in some tumors formed in gels that may represent residual gel material. This material is designated by black arrows. Upon evaluation by a blinded pathologist, these tumors do not appear morphologically different than tumors formed in BME and saline. Columns represent a single tumor, and rows represent a single magnification. 2x magnification the scale bars represent 2 mm. 4x magnification the scale bars represent 1 mm. 10x magnification the scale bars represent 400 um.

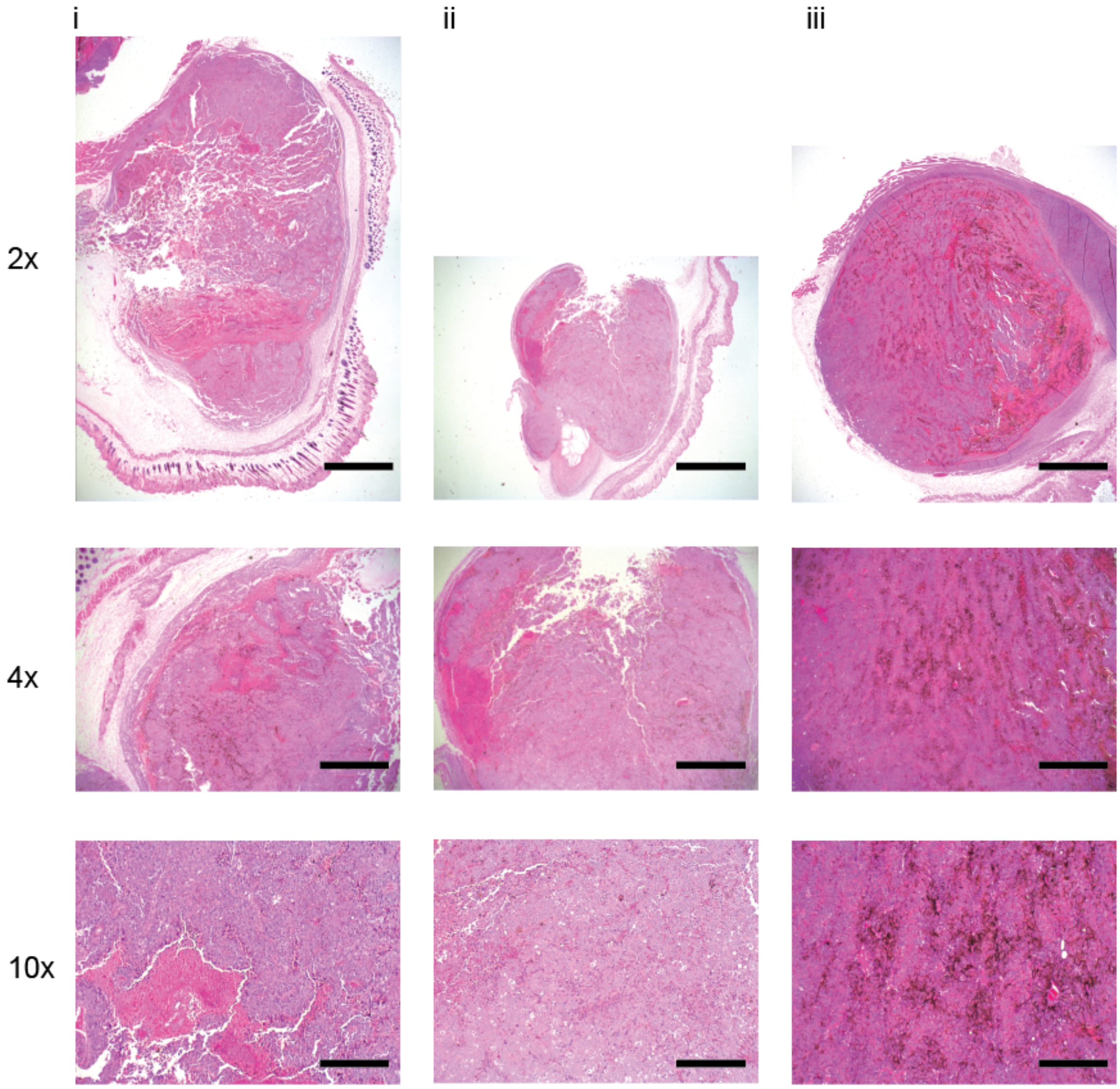


**Supplementary Figure 13:** Additional histology images with Hemotoxylin and Eosin staining from Day 15 for tumors formed in Alginate-Hyaluronic Acid. Upon evaluation by a blinded pathologist, these tumors do not appear morphologically different than tumors formed in BME and saline. Tumors formed in Alg-HA were the largest on averge of all the gel groups. Columns represent a single tumor, and rows represent a single magnification. 2x magnification the scale bars represent 2 mm. 4x magnification the scale bars represent 1 mm. 10x magnification the scale bars represent 400 um.

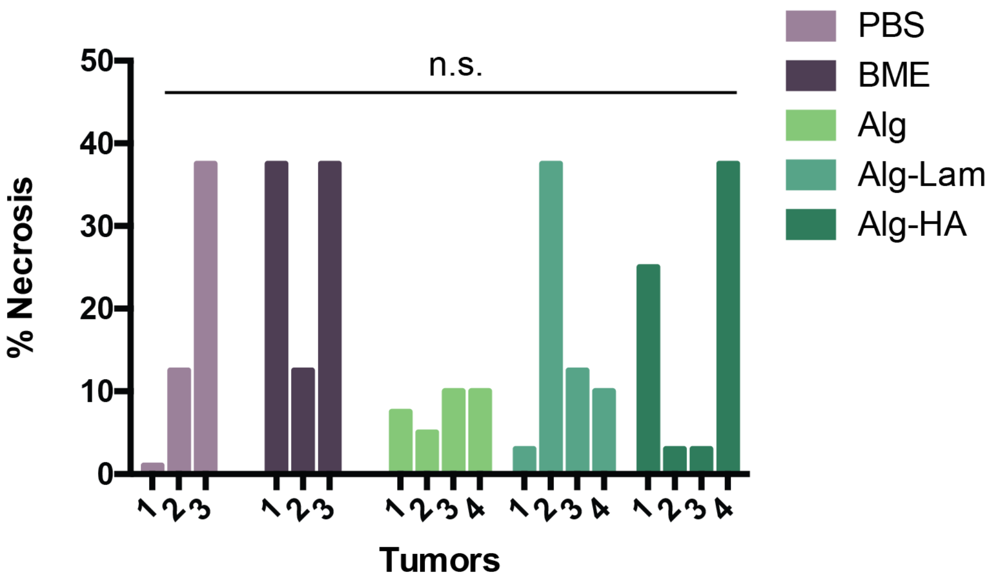


**Supplementary Figure 14:** % Necrosis as determined by a blinded pathologist for each tumor in all experimental groups.

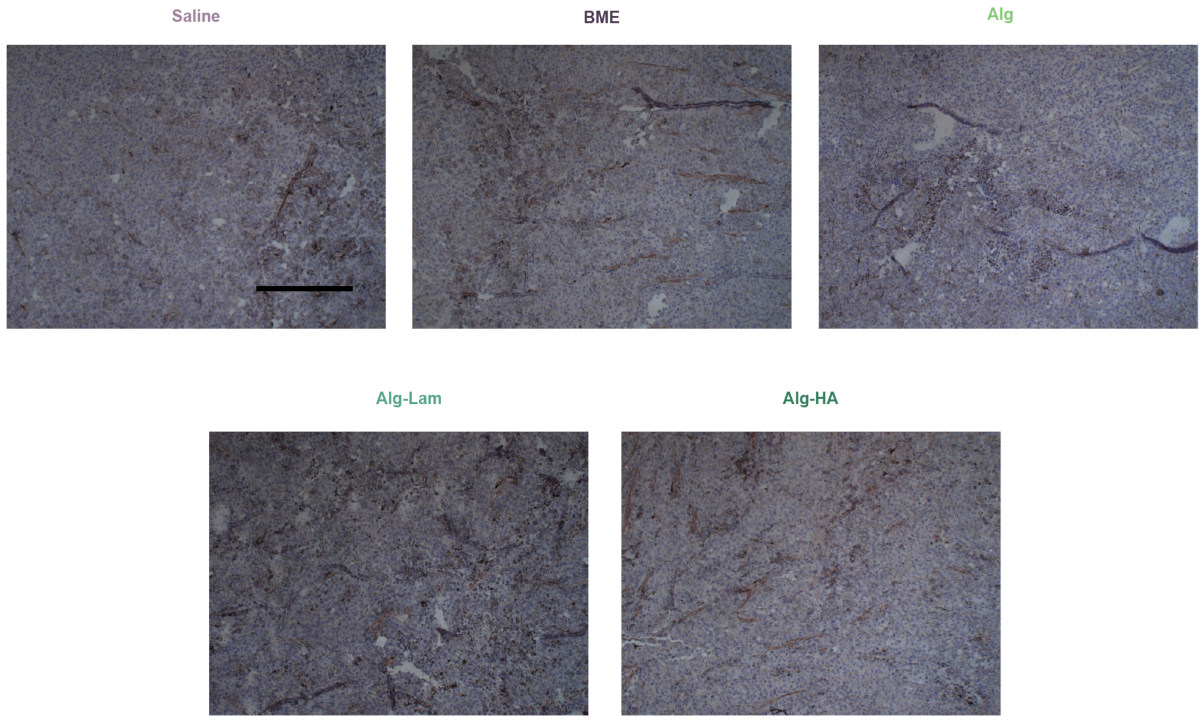


**Supplementary Figure 15:** Representative CD31 staining from each group. No significant differences were observed by a blinded pathologist. Background melanin made clear analysis challenging. Scale bar represents 250 um.

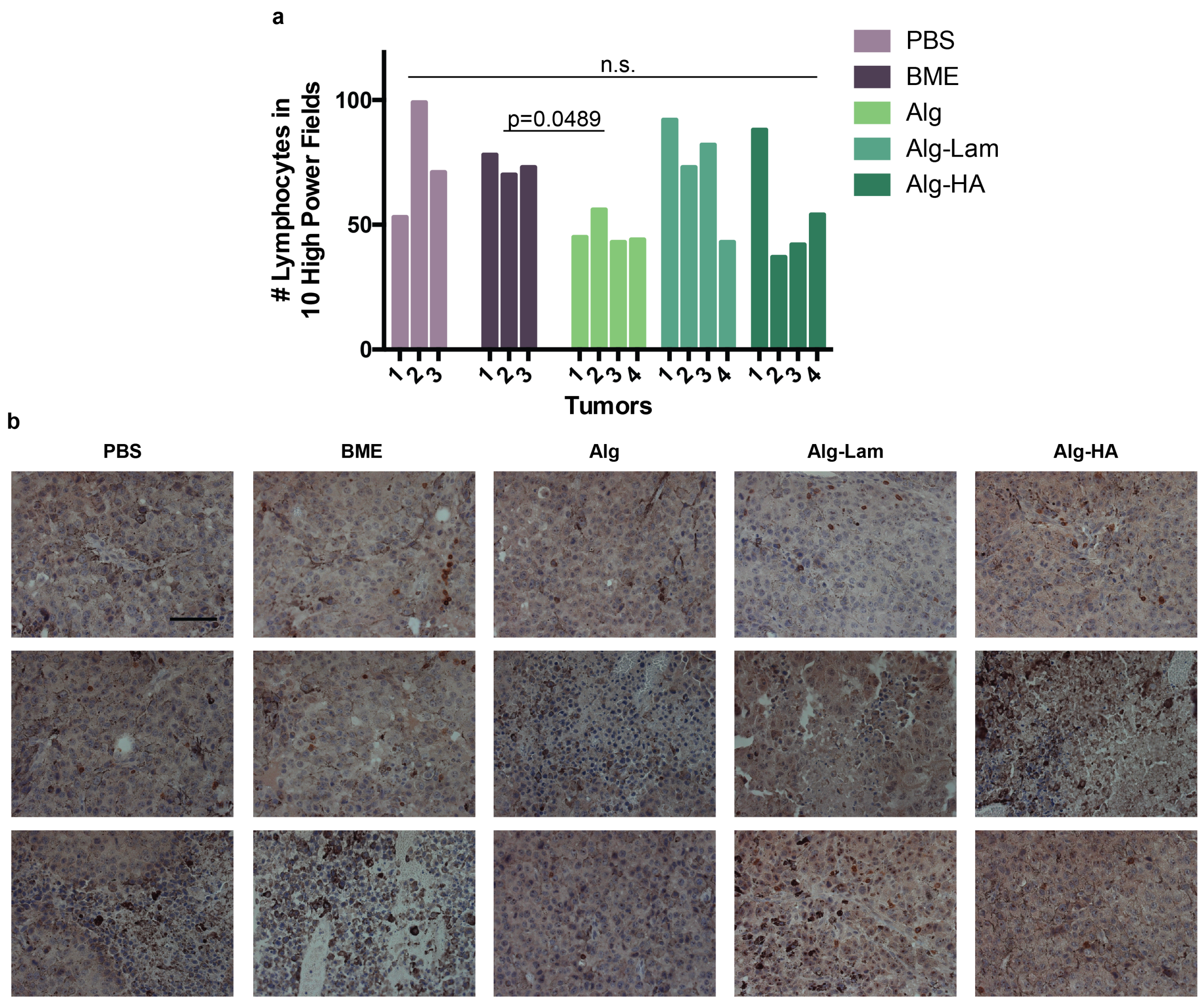


**Supplementary Figure 16:** **a,** Number of lymphocytes per tumor found in 10 high power fields (40x magnification) for all experimental groups. **b,** Representative high power images of CD3 stained tumors where lymphocytes are stained dark brown. Scale bar represents 50 um.

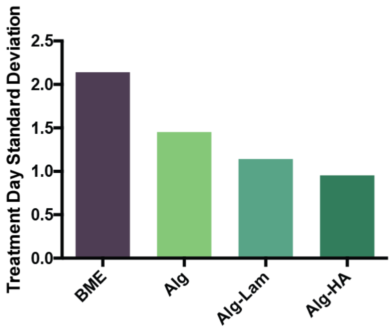


**Supplementary Figure 17:** Standard deviation of treatment day for each formulation. Standard deviations from the averages shown in Fig 3c.

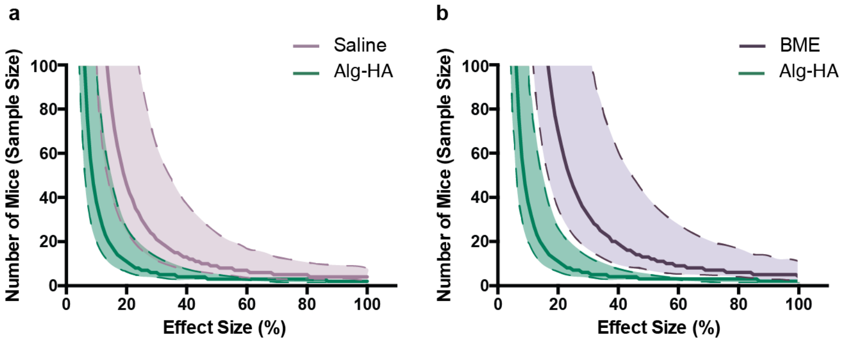


**Supplementary Figure 18: a-b,** Power analysis predicting the number of mice per group needed based off the experimental data obtained with 80% power. The dashed lines and shading represent the high and low number estimate of mice given the high and low 95% confidence interval values on the input coefficient of variance for these power calculations. Confidence intervals on coefficient of variation were calculated using Vangel's modification on McKay's theory on coefficient of variance. The Statistical toolbox in Matlab was used for power calculations, specifically the *sampsizepwr* function with two-sided t tests. The average and distribution of the day that each formulation surpassed 100 $mm^2$ were used in the power analysis.

### Supplementary Methods

##### Sample Matlab code for power analysis calculations

```
%Code for Fig 2i
Power2 = 0.8;
End = 100;
samplesizePBS2 = [];
for x = 0:0.01:End
n = sampsizepwr('t2',[averagePBS_day15 , stdPBS_day15], x*averagePBS_day15, ...
    Power2,[]);
samplesizePBS2 =  [ samplesizePBS2 n];
end
samplesizebme2 = [];
for x = 0:0.01:(End)
n = sampsizepwr('t2',[averagebme_day13 , stdbme_day13], x*averagebme_day13, ...
    Power2,[]);
samplesizebme2 =  [samplesizebme2 n];
end
samplesizehighHA2 = [];
for x = 0:0.01:End
n = sampsizepwr('t2',[averagehighHA_day13 , stdhighHA_day13], ...
    x*averagehighHA_day13, Power2,[]);
samplesizehighHA2 =  [ samplesizehighHA2 n];
end


%Code for Fig 2j
End = 50;
samplesize1 = [];
for x = 1:1:End
n = sampsizepwr('t2',[100, x], 60, Power2,[]);
samplesize1 =  [samplesize1 n];
end
samplesize2 = [];
for x = 1:1:End
n = sampsizepwr('t2',[100, x], 70, Power2,[]);
samplesize2 =  [samplesize2 n];
end
samplesize3 = [];
for x = 1:1:End
n = sampsizepwr('t2',[100, x], 80, Power2,[]);
samplesize3 =  [samplesize3 n];


%Code for Fig 2k
End = 50;
power1 = [];
for x = 1:1:End
pwrout = sampsizepwr('t2',[100, x], 80,[],10);
pwrout = pwrout*100;
power1 =  [power1 pwrout];
end
power2 = [];
for x = 1:1:End
pwrout = sampsizepwr('t2',[100, x], 70,[],10);
pwrout = pwrout*100;
power2 =  [power2 pwrout];
end
power3 = [];
for x = 1:1:End
pwrout = sampsizepwr('t2',[100, x], 60,[],10);
```

```
pwrout = pwrout*100;
power3 =  [power3 pwrout];
end
```

#### Supplementary Videos

**Supplementary Video 1: Formulation of cargo-loaded alginate hydrogels by simple mixing.** The hydrogel is prepared with a dual syringe mixer whereby a 5 wt% polymer solution is loaded in one syringe (right), and cells and 10 mM calcium sulfate are placed in the second syringe (left, dyed blue for this demonstration for easier viewing). The two solutions are mixed with a female-female dual syringe mixer by pumping the material back and forth for 30 seconds to form a robust hydrogel that is already loaded into a syringe and ready for administration.

**Supplementary Video 2: Injectability of cargo-loaded self-assembled hydrogels.** These alginate-based hydrogels are easily injected from standard syringes after mixing and loading into syringes. The mechanical properties of the hydrogel, which is dyed blue for this demonstration for easier viewing, prevent settling of cargo and local retention in a depot following injection.
